# crisprQTL mapping as a genome-wide association framework for cellular genetic screens

**DOI:** 10.1101/314344

**Authors:** Molly Gasperini, Andrew J. Hill, José L. McFaline-Figueroa, Beth Martin, Cole Trapnell, Nadav Ahituv, Jay Shendure

**Author notes:** Correspondence to Jay Shendure and Molly Gasperini.

## Abstract

Expression quantitative trait locus (eQTL) and genome-wide association studies (GWAS) are powerful paradigms for mapping the determinants of gene expression and organismal phenotypes, respectively. However, eQTL mapping and GWAS are limited in scope (to naturally occurring, common genetic variants) and resolution (by linkage disequilibrium). Here, we present crisprQTL mapping, a framework in which large numbers of CRISPR/Cas9 perturbations are introduced to each cell on an isogenic background, followed by single-cell RNA-seq (scRNA-seq). crisprQTL mapping is analogous to conventional human eQTL studies, but with individual humans replaced by individual cells; genetic variants replaced by unique combinations of ‘unlinked’ guide RNA (gRNA)-programmed perturbations per cell; and tissue-level RNA-seq of many individuals replaced by scRNA-seq of many cells. By randomly introducing gRNAs, a single population of cells can be leveraged to test for association between each perturbation and the expression of any potential target gene, analogous to how eQTL studies leverage populations of humans to test millions of genetic variants for associations with expression in a genome-wide manner. However, crisprQTL mapping is neither limited to naturally occurring, common genetic variants nor by linkage disequilibrium. As a proof-of-concept, we applied crisprQTL mapping to evaluate 1,119 candidate enhancers with no strong *a priori* hypothesis as to their target gene(s). Perturbations were made by a nuclease-dead Cas9 (dCas9) tethered to KRAB, and introduced at a mean ‘allele frequency’ of 1.1% into a population of 47,650 profiled human K562 cells (median of 15 gRNAs identified per cell). We tested for differential expression of all genes within 1 megabase (Mb) of each candidate enhancer, effectively evaluating 17,584 potential enhancer-target gene relationships within a single experiment. At an empirical false discovery rate (FDR) of 10%, we identify 128 *cis* crisprQTLs (11%) whose targeting resulted in downregulation of 105 nearby genes. crisprQTLs were strongly enriched for proximity to their target genes (median 34.3 kilobases (Kb)) and the strength of H3K27ac, p300, and lineage-specific transcription factor (TF) ChIP-seq peaks. Our results establish the power of the eQTL mapping paradigm as applied to programmed variation in populations of cells, rather than natural variation in populations of individuals. We anticipate that crisprQTL mapping will facilitate the comprehensive elucidation of the *cis-*regulatory architecture of the human genome.

## Main Text

Consequent to an era of biochemical surveys of the human genome (such as the ENCODE Project, (*1*)) and ‘common variant’ human genetics (*i.e.* GWAS, eQTL, (*2*, *3*)), we are currently awash in candidate regulatory elements and phenotype-linked haplotypes, respectively. It is clear that determining whether and how each candidate regulatory element is truly functional, as well as pinpointing which noncoding variants are causal for genetic associations, will require functional characterization of vast numbers of sequences, *e.g.* through massively parallel reporter assays and/or CRISPR/Cas9-directed perturbations.

We and other groups have recently adapted cell-based CRISPR/Cas9 genetic screens to evaluate candidate regulatory sequences in their native genomic context ((*4*–*12*)). However, two key aspects of the methods used in these studies limit their scalability. First, each study focuses on the regulation of a single gene per experiment, typically entailing the development of a gene-specific assay. Second, each cell is a vehicle for a single perturbation, *e.g.* CRISPR-mediated repression, mutation or deletion of one candidate regulatory sequence per cell, with the specificity-conferring gRNAs usually introduced via lentivirus at a low multiplicity of infection (MOI). With millions of candidate regulatory elements and ~20,000 regulated genes in the human genome, these limitations preclude the comprehensive dissection of the *cis-*regulatory architecture even within a single cell line.

crisprQTL mapping (**Fig. 1A**) is designed to overcome both limitations. First, by using scRNA-seq instead of gene-specific assays, a single experiment can globally capture perturbations to gene expression, with no strong *a priori* hypothesis as to the target gene of each regulatory element tested. Second, by introducing gRNAs at a high MOI, each individual cell acquires a unique combination of ‘unlinked’ perturbations against the isogenic background of a cell line. Introducing multiple perturbations per cell greatly increases the power to detect changes in gene expression and reduces the number of cells required per experiment (**Fig. 1B**). As in human eQTL studies (*13*, *14*), an association framework can be used to map *cis-* and *trans-*effects by comparing gene expression in the subsets of cells that contain each gRNA to the cells that lack that guide (rather than comparing the subsets of people that do vs. don’t harbor each genetic variant). However, unlike eQTL studies, the resolution of crisprQTL mapping is not constrained by linkage disequilibrium, nor is it limited to studying the effects of naturally occurring, common genetic variants.

**Fig. 1.**
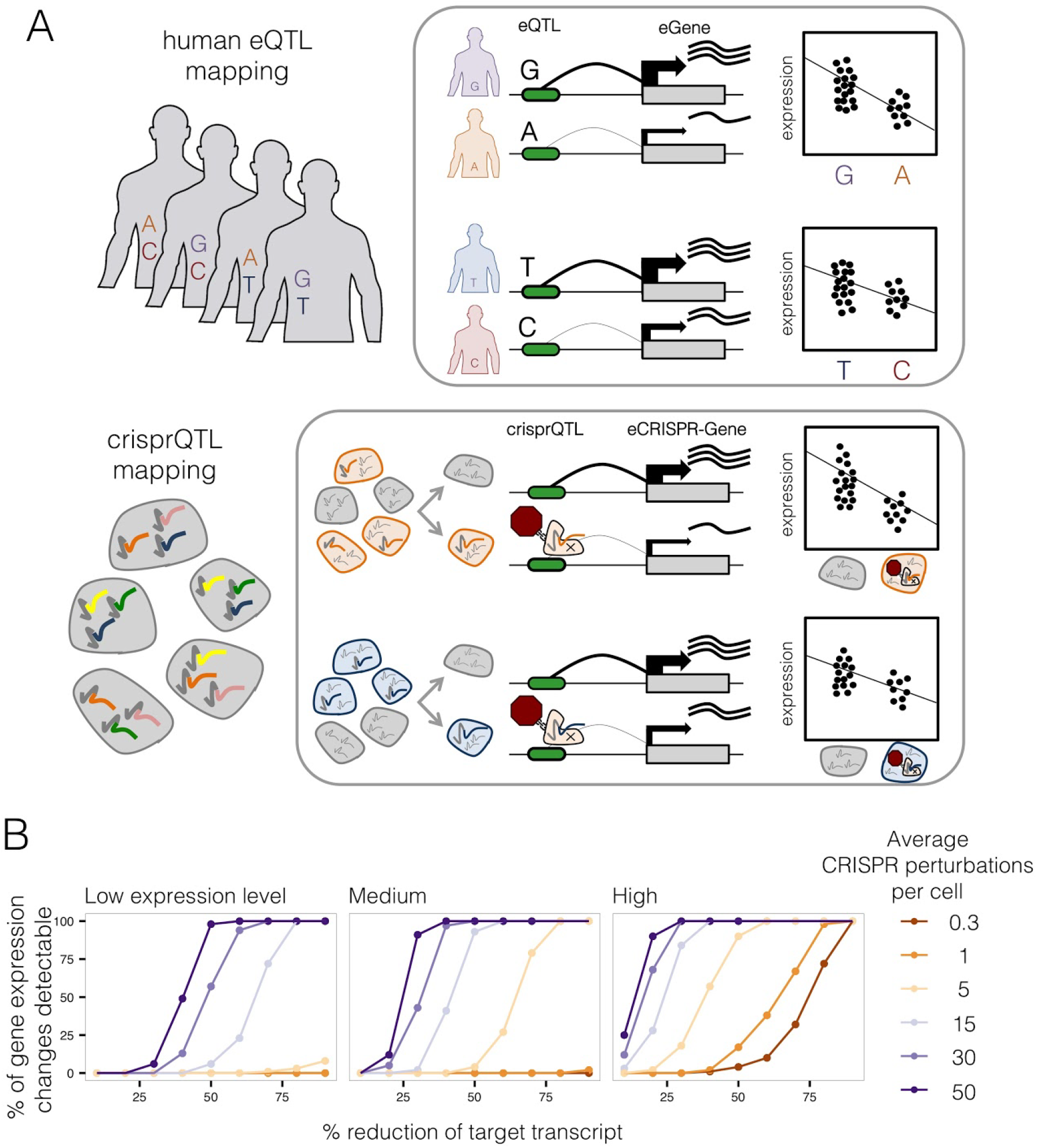
crisprQTL mapping. (A)crisprQTL mapping uses the same framework as human eQTL studies, but with a population of human individuals replaced by a population of individual cells, natural genetic variation replaced by diverse combinations of gRNA-programmed perturbations in each cell, and tissue-level RNA-seq of each person replaced by scRNA-seq. **(B)** Multiplex perturbations increase power to detect changes in gene expression in single-cell genetic screens while greatly reducing the number of cells necessary to profile. Simulated power calculations show that increasing the average number of perturbations per cell (*e.g.*, by increasing MOI in lentiviral delivery of gRNAs) strongly increases power to detect changes in gene expression, including for genes with low (0.10 mean UMIs per cell), medium (0.32) or high (1.00) levels of mean expression. X-axis corresponds to the simulated % change of transcript repressed by targeting CRISPRi to the associated enhancer. Calculations assume a fixed number of 45,000 cells profiled by scRNA-seq.

As an initial pilot of crisprQTL mapping, we targeted 100 candidate enhancers in the chronic myelogenous leukemia cell line K562, with CRISPR-interference (CRISPRi) as our mode of perturbation. For CRISPRi, we used a nuclease-inactive Cas9 tethered to the KRAB repressor domain to induce heterochromatin across a 1-2 kilobase (Kb) window around a gRNA’s target site (*15*). The 100 candidate enhancers were all intergenic sites of coincident DNase I hypersensitivity (DHS), H3K27 acetylation and GATA1 binding, that resided within the same topologically associated domain (TAD) as at least one gene with high K562-specific expression (**Fig. 2A**). Two gRNAs were designed to target each candidate enhancer. Additional pairs of gRNAs served as positive (targeting the transcription start sites (TSSs) of 25 highly expressed genes, or hypersensitivity sites of the *β*-globin locus control region (LCR)) or negative controls (13 non-targeting controls or ‘NTC’ that either fall nowhere in the genome or in a gene desert) (**Table S1**).

This pilot gRNA library was cloned into the lentiviral CROP-seq vector modified to include a CRISPRi-optimized backbone (*16*, *17*) and K562 cells were transduced at a high MOI (**Fig. 2B**). After 10 days to allow for effective CRISPRi, the transcriptomes of 6,806 single cells were sequenced on the 10x Genomics platform (**Fig. 2C**). With a targeted enrichment protocol (*18*), we identified a median of 8 +/- 4.5 gRNAs per cell (interquartile range = 7, **Fig. S1A**). The mean ‘allele frequency’ of perturbations in this experiment – that is, the proportion of cells in which a given candidate enhancer or control was targeted – was 4.1% +/- 2.2% (**Fig. S1B**). For each targeted element, we partitioned the 6,806 cells based on whether they did or did not contain gRNA(s) targeting it (**Fig. 2D**). We then tested for reductions in expression of all genes within 1 Mb of that element (**Fig. 2E**), similar to *cis* eQTL identification in conventional eQTL studies (*14*).

The 13 NTC controls were tested against all 1,196 genes that fell within 1 Mb of any targeted element (**Fig. 2F**). We defined a 10% empirical FDR threshold based on these NTC tests as they are subject to the same sources of error as the element-targeting gRNAs. With this threshold, 68% (17 of 25) of TSS-targeting positive controls repressed their associated genes, as did the *β*-globin LCR controls (**Fig. 2G**). In contrast, targeting of only 3% of the candidate enhancers (3 of 100) significantly decreased expression of any gene within 1 Mb (**Fig. 2F**, **Fig. S1C**). All three decreased expression of the same gene, *NMU*, which encodes neuromedin U, a neuropeptide that plays roles in inflammation as well as erythropoiesis ((*19*–*22*)). These three crisprQTLs were relatively closely located: 93.4, 94.1 and 97.6 Kb upstream of the *NMU* TSS (ii-iv in **Fig. 2H**; region denoted ‘e-NMU’ as potentially too close to one another to resolve by CRISPRi). In contrast, targeting of a different candidate enhancer located just 34.4 Kb upstream of the *NMU* TSS (i in **Fig. 2H**) did not significantly reduce its expression.

**Fig. 2.**
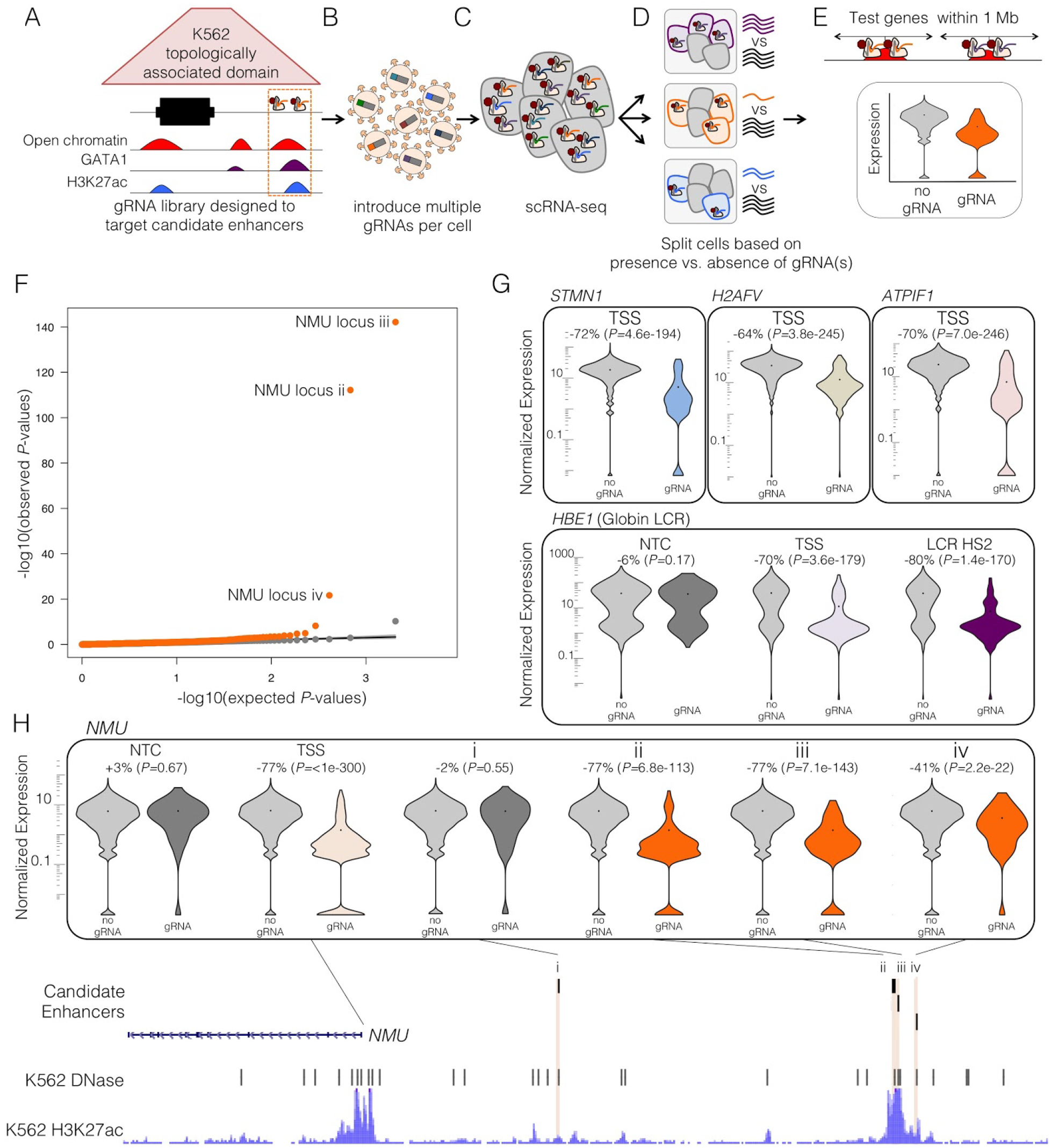
Pilot crisprQTL mapping study testing 100 candidate enhancers in K562 cells. (A-E), Schematic of crisprQTL mapping. **A,** Candidate enhancers were chosen based on enhancer-associated features, and were each targeted by 2 gRNAs. **B,** gRNAs were cloned into the CRISPRi-optimized CROP-seq lentiviral vector, and delivered to K562 cells at a high MOI. **C,** scRNA-seq was performed on these cells, with concurrent identification of the combination of gRNAs present in each cell. **D,** For each candidate enhancer, cells were partitioned based on whether or not they contained a gRNA targeting it. **E,** For each such partition, we used Monocle (*23*) to test for differential expression between the two populations for any gene within 1 Mb of the candidate enhancer. **(F)** Quantile-quantile (Q-Q) plot of the differential expression tests. Distributions of observed vs. expected *P*-values for candidate enhancer-targeting gRNAs (orange) and NTC gRNAs (gray; downsampled) are shown. **(G)** Expression of selected TSS and, *β*-globin LCR positive controls. Positive controls showed significant differential expression of the expected target genes between cells *with* versus *without* targeting gRNAs, in contrast with NTCs. **(H)** Distal enhancers of *NMU* identified as crisprQTLs. Targeting three candidate enhancers (labeled *ii-iv*) located 93.4, 94.1 and 97.6 Kb upstream of *NMU*, significantly reduced expression of *NMU*, but targeting an intervening candidate enhancer (labeled *i*) located 34.4 Kb upstream did not.

To demonstrate crisprQTL mapping at scale, we designed gRNAs targeting 1,119 candidate enhancers throughout the genome (2 gRNAs each; 62/100 elements from pilot experiment were re-targeted, including the three crisprQTLs upstream of *NMU*). For this experiment, we adjusted criteria for selection, focusing on intergenic DHS that exhibited various combinations of H3K27ac, p300 binding, GATA1 binding and RNA Pol II binding (**Fig. 3A**). Candidate enhancers were required to fall within the same TAD (*24*) as at least one gene exhibiting strong K562-specific expression (top 10% most highly expressed genes in the 6,806 cells of the pilot dataset), and were collectively distributed across 510 TADs on every chromosome. Almost half of all genes that are expressed in K562 cells fell within 1 Mb of at least one targeted candidate enhancer (5,751 of the 12,984 expressed genes; expressed = observed in at least 0.525% of cells in this experiment). In addition, pairs of gRNAs targeting 381 TSSs (of genes sampled from the top decile of K562 expression), the globin LCR and 50 NTCs were again included as positive or negative controls (**Table S2**).

We cloned these gRNAs into the CRISPRi-optimized CROP-seq vector, transduced K562 cells at a high MOI, and then generated 47,650 scRNA-seq profiles. With greater input material to the targeted amplification than was used in the pilot study, we identified a median of 15 +/- 11.3 gRNAs per cell (interquartile range 15, **Fig. 3B**). The mean ‘allele frequency’ of perturbations in this experiment was 1.1% +/- 0.06% (**Fig. 3C**). We then iteratively partitioned the cells based on whether they did or did not contain gRNA(s) targeting each TSS or candidate enhancer, and tested for reductions in expression of all genes within 1 Mb of it (**Fig. S2**). We also tested the 50 NTCs against all 5,751 genes within 1 Mb of any targeted candidate enhancer. For perspective, in a ‘one gRNA per cell’ framework, achieving equivalent power would require generating ~715,000 single cell transcriptomes.

**Fig. 3.**
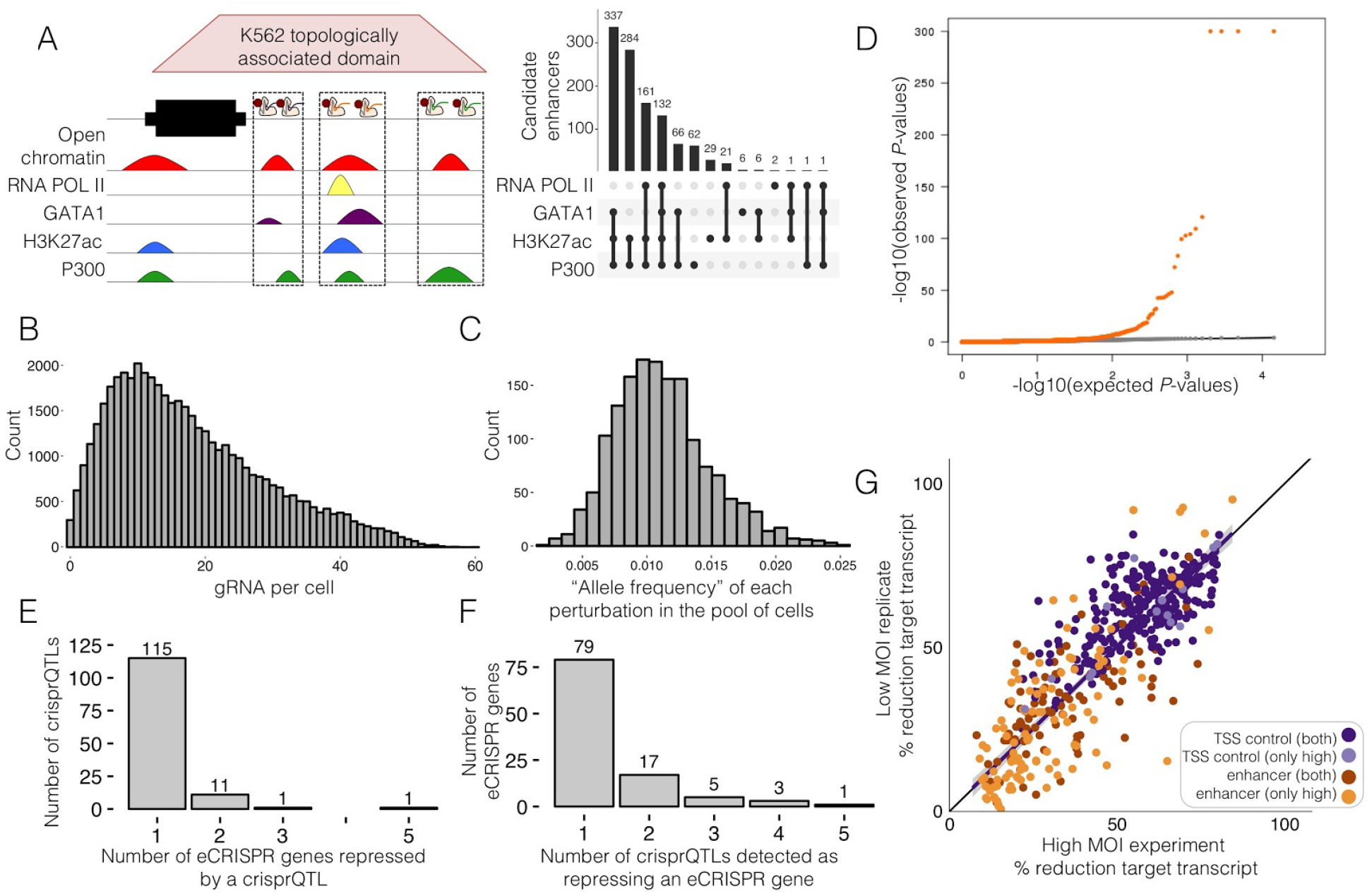
crisprQTL mapping at scale in K562 cells. (A)gRNAs were designed to target 1,119 candidate enhancers, as defined by DNase hypersensitive sites intersecting with 1+ enhancer-associated biochemical marks. **(B)** gRNAs were delivered to K562 cells at a high MOI (median 15 +/- 11.3 gRNAs identified per cell). **(C)** The median ‘allele frequency’, *i.e.* the proportion of the 47,650 cells in which a given element was targeted, was 1.1% +/- 0.06%. **(D)** Q-Q plot of the differential expression tests. Distributions of observed vs. expected *P*-values for candidate enhancer-targeting gRNAs (orange) and NTC gRNAs (gray; downsampled) are shown. **(E)** Histogram of the number of eCRISPR-genes significantly impacted by crisprQTLs. **(F)** Histogram of the number of crisprQTLs detected as regulating an eCRISPR-gene. **(G)** Comparison of effect sizes from the original high MOI study and a low MOI replicate (41,284 cells, median 1 gRNA/cell). Only TSS or candidate enhancers that met thresholds in the high MOI study are plotted (Spearman’s rho 0.826; slope 0.9995, y-intercept −0.011; both = identified in both low and high MOI study; only high = only in the high MOI study).

A quantile-quantile plot showed an excess of significant associations involving the targeting of candidate enhancers, relative to NTC controls (**Fig. 3D**). Setting a threshold corresponding to a 10% empirical FDR (based on the NTCs), 94% (357 of 381) of TSS-targeting positive controls repressed their associated genes, as did the *β*-globin LCR controls. In addition, targeting of 11% (128 of the 1,119) of candidate enhancers repressed 1+ gene(s) within 1 Mb. As there were 13 candidate enhancers whose targeting impacted more than one gene (**Fig. 3E**), our data identified a total of 145 crisprQTL enhancer-gene relationships (**Table S3**). Of the 105 downregulated genes (‘eCRISPR genes’), 26 were impacted by targeting of more than one candidate enhancer in the library (**Fig. 3F**).

The three candidate enhancers implicated as crisprQTLs of *NMU* in the pilot experiment were again identified here, as was an enhancer 3.6 Kb upstream of *GATA1* that was previously found in a tiling CRISPRi-noncoding screen (*11*). Additional examples are highlighted in **Fig. 4**. Our data support three elements as crisprQTLs of *FEZ1*, a gene implicated in schizophrenia (*25*, *26*) that is nonetheless expressed in K562 cells (**Fig. 4A**). One falls 37 Kb upstream (i; **Fig. 4A**), while the other two are clustered 56 Kb and 57 Kb upstream (ii-iii; **Fig. 4A**). Targeting these latter two elements also decreased expression of nearby *EI24*, which is 17 Kb away in the opposite direction. Our data also support a crisprQTL within the fatty acid desaturase (*FADS*) cluster. The *FADS* locus is thought to have experienced multiple independent episodes of local adaptation in geographically diverse human populations in response to differences in diet ((*27*–*29*)). The candidate enhancer lies between *FADS2* and *FADS3* (which are oriented tail-to-tail), but downregulation is only detected for the more distal *FADS1* (**Fig. 4B**).

A longstanding question in gene regulation is by what distances are endogenous enhancers separated from their target genes in the human genome? We find crisprQTLs were separated from the TSS of their target genes by a median distance of 34.3 Kb (upstream crisprQTLs only, to avoid biasing of this calculation by our avoidance of intronic candidate enhancers), supporting a strong role for proximity in governing enhancer-promoter choice, given that we tested association of all genes within 1 Mb (**Fig. 4C**). However, the majority of crisprQTL regulatory relationships did not involve the most proximal gene, and not even the most proximal K562-expressed gene (**Fig. 4D**; only 34/145 (23%) of the crisprQTLs detected in this experiment involved the K562-expressed gene most proximal to the targeted candidate enhancer).

Of our 357 “positive control” TSSs whose targeting successfully repressed the expected gene, 55 (15%) reduced expression of 1+ additional genes (72 apparent promoter-promoter relationships in total). 16 of these 72 involved overlapping promoters (TSSs within 1 Kb), such that the observed effect of CRISPRi is likely direct. As for the remaining 56, one possibility is that these represent examples of promoters acting as enhancers, as has been recently reported (*11*, *12*). However, these 56 are *not* enriched for proximity to affected genes (median distance of upstream genes = 448.6 Kb; median distance to all genes within 1 Mb = 441 Kb) (**Fig. S3**), in contrast with the 145 enhancer-TSS crisprQTL relationships (median distance = 34.3 Kb) (**Fig. 4C**). We therefore hypothesize that these are more likely *trans* effects of repressing the primary target.

The set of 105 eCRISPR genes identified here, *i.e.* genes regulated by 1+ crisprQTL(s), had several notable characteristics. First, their expression levels largely fall in the top half of all 5,751 genes against which we tested (**Fig. 4E**), suggesting that we are likely underpowered to detect regulatory effects on more lowly expressed genes. Second, we detected no depletion of housekeeping genes (hypergeometric test; *P*-value = 0.95, 34 of 105 (32%) genes annotated as housekeeping (*30*) vs. 35% of all genes expressed in K562), arguing against the view that a prevailing characteristic of housekeeping genes may be a relative dearth of distal regulatory elements (*5*, *31*). Finally, functional annotation enrichment of the 105 eCRISPR genes was most significant for hematopoiesis and regulation of erythrocyte differentiation, consistent with distal enhancers primarily shaping the expression of K562 cell type-specific genes (**Table S4**) (*32*, *33*). These terms were not enriched in a random expression-level matched control gene set.

We also examined the characteristics of crisprQTLs whose targeting significantly impacted expression of 1+ genes in *cis.* Among the 1,119 total candidate enhancers targeted, the 128 crisprQTLs were positively correlated with ChIP-seq peak strength (based on average enrichment in ChIP-seq peak region) for enhancer-associated histone modifications (H3K27ac, logistic regression *P*-value = 4e-05), co-activators (p300, *P*-value = 4e-16) and lineage-specific TFs (GATA1 *P*-value *=* 2e-07, GATA2 *P*-value = 3e-10, SMAD1 *P*-value = 1e-06, TAL1 *P*-value = 6e-06, CCNT2 *P*-value = 3e-07), whereas RNA Pol II and H3K4me1 were not associated (**Fig. 4F**). Because many of these biochemical features are correlated, we trained a multivariate logistic regression classifier to ask whether we could distinguish the 128 crisprQTLs from the 991 candidate enhancers for which we did not identify an eCRISPR target gene, achieving an AUPR of 0.31 (area under precision-recall curve; median from five-fold cross validation, **Fig. S4**). H3K27ac, p300 and distance from the closest TSS were the only significant predictors in the multivariate model (**Table S5**).

To replicate a small subset of our crisprQTLs outside of the multiplexed screen, we separately transduced small pools of gRNA that targeted individual crisprQTL candidate enhancers, and investigated repression of eCRISPR gene expression levels via bulk RNA-seq. For such experiments targeting eNMU, ePRKCB, ePTGER3, and eGYPC, we observed effect sizes on the expected eCRISPR target gene that matched the effect size from the pooled mapping experiment (crisprQTL repression ranging from −79% to −42%, *P*-values from 6.8e-113 to 6.2e-11, **Fig. S5**). For these four cases, the expected eCRISPR gene was the top differentially expressed target gene. Although eHMGA1 failed to replicate, we note that of the five, it was the crisprQTL closest to our 10% empirical FDR threshold, with only −11% repression and *P*-value of 0.00033.

**Fig. 4.**
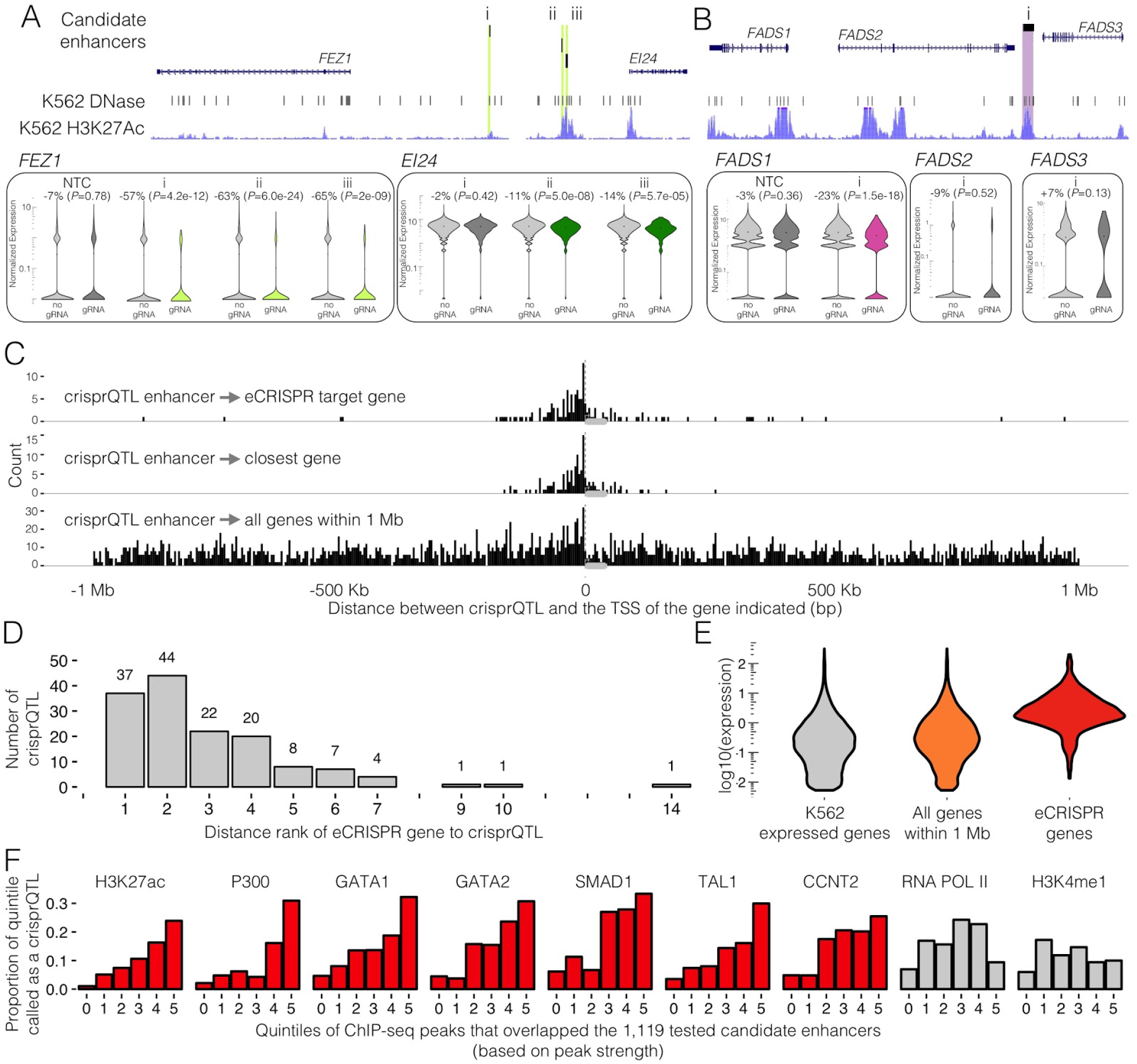
Examples and characteristics of K562 crisprQTLs. **(A)** crisprQTLs of *EI24* and *FEZ1*. When targeted in our screen, three candidate enhancers (i-iii) decreased expression of the schizophrenia-associated gene *FEZ1* (*25*), with the latter two (ii-iii) also decreasing nearby *EI24*, a putative tumor suppressor (*34*) (shown: chr11:125,363,009-125,443,355, hg19). **(B)** Insight into regulation of the *FADS* locus, a region under positive selection in geographically-diverse human populations. A candidate enhancer (i) lies between the fatty acid desaturases *FADS2* and *FADS3*, which are oriented tail-to-tail, but targeting it only decreased expression of the neighboring *FADS1* (shown: chr11:61,566,537-61,660,119, hg19). **(C)** crisprQTLs fall close to target genes. Distribution of distances between the crisprQTLs’ candidate enhancers and: their target eCRISPR gene’s TSS (top), the TSS of whatever K562-expressed gene is closest (middle), or the TSS of any gene within 1 Mb (bottom). Plotted with respect to gene orientation. **Fig. S8** presents a zoomed in view of +/- 20 Kb from the TSS. Of note, candidate enhancers within 1 Kb of any gene body, *i.e.* including all exonic or intronic candidate enhancers, were excluded in library design. **(D)** 111 of 148 crisprQTLs do not target the most proximal K562-expressed gene. Genes are ranked by their distance to the candidate enhancer (1 = closest, 2 = second closest, *etc.*). **(E)** crisprQTL mapping is better powered to detect regulatory effects on highly expressed genes (expression = mean reads/cell in the entire 47,650 cell dataset). **(F)** crisprQTLs tend to fall in enhancer-associated ChIP-seq peaks that show stronger signals. All ChIP-seq peaks that overlap the 1,119 candidate enhancers were divided into quintiles of strength, defined as the average enrichment in ChIP-seq peak region (0 = no such peak overlaps the candidate enhancer, 1 = lowest, 5 = highest). Histograms of the proportion of each 1,119-quintile that were called as crisprQTLs are shown. Red = *P*-value < 0.005 for independent logistic regression for predicting a candidate enhancer as a crisprQTL based on this peak type.

One concern was whether multiple gRNAs within a single cell would dilute the efficacy of CRISPRi. To evaluate this, we conducted a biological replicate experiment with the same gRNA library but at a low MOI (41,284 cells, median 1 gRNA/cell, standard deviation=1.6, interquartile range = 1, **Fig. S6A**). The effect sizes identified in the high MOI experiment were well correlated with the effect sizes for the same gRNAs in the low MOI experiment (Spearman’s rho: 0.826, slope: 0.9995, **Fig. 3G**). However, at the same threshold corresponding to a 10% empirical FDR, only 265 TSSs and 20 crisprQTLs were identified in the low MOI experiment (**Fig. S6B-D**), validating the substantial increase in power resulting from our multiplexed perturbation framework (**Fig. 1B**).

Even though this experiment solely targeted regulatory elements with CRISPRi, *trans* effects are still possible, *e.g.* as the secondary consequences of *cis* downregulation of a transcription factor. We therefore repeated our association testing, but evaluated each perturbation (TSS or candidate enhancer) against all K562-expressed genes located >1 Mb from the target gene or on another chromosome. Once again setting a threshold at an empirical FDR of 10% based on equivalent tests on NTCs, we identified 15 *cis* crisprQTLs that are also associated with downregulation of 1+ genes >1 Mb away or on other chromosomes (**Fig. S7A**). In addition, targeting 130 of the TSSs (targeting of which we previously associated with downregulation of the cognate gene) also resulted in apparent *trans* effects, with 81 (62%) of these influencing >1 distally located gene (**Fig. S7A**). For example, targeting the TSS of *MRPL33* successfully repressed both *MRPL33* and 112 other genes. As *MRPL33* encodes the 39S large subunit of mitochondrial ribosomes, affected genes were correspondingly most enriched for electron transport chain and mitochondrial pathways (strongest ten *q*-values).

Understanding the regulatory landscape of the human genome requires that we validate and identify the target genes of vast numbers of candidate enhancers, as nominated by biochemical marks and eQTL mapping. crisprQTL mapping has the potential to meet this challenge. In a single experiment, we evaluated 17,584 potential *cis* regulatory relationships involving 1,119 candidate enhancers and 5,751 expressed genes. In contrast, nine recently published CRISPR screens of noncoding sequences cumulatively studied regulatory effects on a total of 17 genes ((*4*–*12*)). By delivering a median of 15 perturbations to each of 47,650 cells, this experiment was powered equivalently to a ‘one gRNA per cell’ experiment involving 715,000 single cell transcriptomes. Such an experiment would be amongst the largest scRNA-seq datasets generated to date and therefore our framework represents enormous savings in cost and labor. Of note, one recent study used scRNA-seq as a readout for the effects of CRISPR-based perturbations of 71 candidate regulatory elements on ~100 genes in seven genomic regions (*35*). However, its power and scope was limited by a low MOI (**Fig. 1B**) and a gRNA barcoding strategy that suffers from a ~50% rate of template switching (*18*).

Although it may be surprising that *cis* changes in gene expression were identified for only 11% of the 1,119 candidate enhancers tested, there are several potential caveats to bear in mind: a) Not all enhancers may be susceptible to CRISPRi-KRAB perturbation; b) gRNAs may be variably effective in targeting enhancers; c) We are underpowered to detect effects on lowly expressed genes; d) Shadow enhancers or other redundancies might mask some effects; e) Some enhancers might be required for the initial establishment rather than maintenance of gene expression; f) We did not comprehensively survey the noncoding landscape surrounding each gene, and the marks we used to define candidate enhancers may be excluding some classes of distal regulatory elements. These caveats are respectively addressable in the future by using other epigenetic modifiers or nuclease-active Cas9, by using more gRNAs per candidate enhancer, by performing better powered experiments via more scalable scRNA-seq methods (*36*), by combinatorial perturbation of selected loci (*35*), by using cell models of differentiation, and by densely tiling selected loci with perturbations.

However, the fact that our crisprQTLs are predicted by the strength of enhancer-associated marks (*e.g.* H3K27ac, p300) supports the assertion that we are identifying *bona fide* enhancers, and simultaneously weakens the case for elements that were negative. Also, our study provides relatively unbiased insight into key properties of mammalian enhancers, *e.g.* the distribution of distances between enhancers and their target genes. A full understanding of the precise rules governing enhancer-promoter choice is a topic of great interest, and will be facilitated by the identification of more crisprQTLs.

A limitation of crisprQTL mapping as implemented here relates to the resolution of CRISPRi (on the order of a 1-2 Kb distribution of heterochromatin silencing as induced by KRAB (*15*)). In the future, this can potentially be improved upon by adapting crisprQTL mapping to use single or pairs of gRNAs with nuclease-active Cas9 to disrupt or delete candidate enhancers at the sequence level. A separate concern is whether high MOI transduction is inducing a cellular inflammatory response, and therefore biasing discovery. However, although some genes with roles in inflammation are amongst our eCRISPR genes (*e.g. NMU, IL6*), we did not observe pathway-level enrichment. Moreover, effect sizes in our high MOI and low MOI experiments were very well-correlated.

A key strength of crisprQTL mapping over conventional eQTL studies is that our framework is not limited by natural genetic variants nor by linkage disequilibrium. However, a weakness is that we are only achieving tens of perturbations in each cell, as opposed to the millions of genetic variants in each human. What are the limits to greater multiplexing of CRISPR-mediated perturbations? It will likely be possible to push the effective MOI per cell higher than we have have here, *e.g.* by repeated rounds of transduction. Coupled with further engineering (*e.g.* multiple gRNAs per lentivirus, or alternative delivery systems), it may be possible to introduce hundreds of ‘unlinked’ perturbations per cell. The other avenue to boost power is to simply profile more cells, which may be facilitated by adapting crisprQTL mapping to more scalable single cell transcriptome profiling schemes such as combinatorial indexing (*36*).

Overall, we anticipate that crisprQTL mapping will enable the validation and target gene discovery of large numbers of candidate enhancers in cell lines and primary tissues. It may also facilitate the fine mapping of causal regulatory variants and their target genes within haplotypes implicated in common disease. To date, ENCODE has catalogued 1.3 million candidate regulatory elements based on biochemical marks (http://screen.umassmed.edu/). Although directly testing all of these elements might not be practical in the near term, the characteristics of enhancers validated here will inform predictive model-driven selection of additional candidates to enrich for crisprQTLs in future screens. Such iterative testing and prediction may be critical for fully elucidating *cis-*regulatory architecture in diverse cell types and throughout the human genome.

## Acknowledgements

We thank the entire Shendure, Trapnell, and Ahituv labs, in particular Darren Cusanovich, Anh Leith, Dana Jackson, Gregory Findlay, Lea Starita, Delasa Aghamirzaie, Aaron McKenna, Malte Spielmann, Vikram Agarwal, Jason Klein, Seungsoo Kim, Xiaojie Qui, Jonathan Packer, Rajiv McCoy, Jacob Tome, Silvia Domcke, Fumitaka Inoue, Orry Elor, and Nadja Makki. We additionally thank James Ousey, Kyuho Han, Michael Bassik, Martin Wohlfahrt, Madison Norton, the staff of CustomArray, Inc. and Martin Kircher. This work was supported by awards from the NIH (UM1HG009408 to NA and JS and DP1HG007811 to JS) and training awards from the National Science Foundation and National Institutes of Health (Graduate Research Fellowship and 5T32HG000035 to MG). JS is an Investigator of the Howard Hughes Medical Institute.

## Materials and Methods

### Cell Lines and Culture

K562 cells expressing dCas9-BFP-KRAB (Addgene #46911) were a gift of the Bassik lab, and cultured in RPMI 1640 + L-Glutamine (Gibco), supplemented with 10% fetal bovine serum (Rocky Mountain Biologicals) and 1% penicillin-streptomycin (GIBCO).

### gRNA-library design

#### Note about terminology

A gRNA-group is defined as all the gRNA that are targeting the same candidate enhancer or positive control site (usually 2). To note, all novel TSS and candidate enhancer targeting gRNA-groups are referred to as “perturbative gRNA-groups”, whereas all others are referred to as “control gRNA-groups. The 100 candidate enhancer pilot library will be referred to as the “100plex library”, and the 1,119 candidate enhancer at-scale library will be referred to as the “1,119plex library”.

### 100 Candidate Enhancer Pilot Library (100plex library, **Table S1**)

#### Picking candidate enhancers

Regions of open chromatin were defined from K562 DNase-seq narrowPeaks (ENCSR000EKS) more than 1 Kb away from any gene (GENCODE March 2017 v26lift37). Using bedtools, these regions were intersected with K562 GATA1 ChIP-seq narrowPeaks (ENCSR000EFT, lifted to hg19), H3K27ac ChIP-seq narrowPeaks (ENCSR000AKP, lifted to hg19), and K562 HiC domains (*24*) that contained at least 1 gene in the top 10% FPKM of K562 most expressed genes that were also more highly expressed in K562 (ENCSR000EYO) than in H1-hESC (ENCSR000EYP) or HepG2 (ENCSR000EYR) as called by Cuffdiff (*q-*value < 0.05)(*37*) but were not K562 essential genes (*38*).

#### TSS positive control gRNA

Each of the ‘top expressed’ K562 genes in the utilized HiC domains were targeted by two TSS-proximal gRNA, from protospacers as best scored by the empirical and predicted scores of the hCRISPRiv2 library (*39*). To note, in further analysis, though its TSS was targeted *‘TOMM6’* was excluded from analysis as it was not detected in the K562 scRNA-seq data, though it was in the bulk RNA-seq dataset (ENCSR000EYO).

#### Candidate enhancer gRNA

NGG-protospacers within these candidate enhancers were scored using default parameters of FlashFry (*40*), and the two top quality-scoring per region were used as spacers in the 100plex gRNA library (scores prioritized by Doench2014OnTarget > Hsu2013 > Doench2016CDFScore > otCount, two gRNA per gRNA-group).

#### Non-targeting control (NTC) gRNA

11 scrambled-sequence spacers with no targets in the genome and 11 protospacers targeting 6 gene-devoid regions of the genome (hg19 chr4:25697737-25700237, chr5:12539119-12541619, chr6:23837183-23839683, chr8:11072736-11075236, chr8:23768553-23771053, chr9:41022164-41024664) were chosen as evaluated by Benchling’s CRISPR tool. These were randomly paired to create a gRNA-group. Additionally, a gRNA targeting the CAG promoter was included for a total of 13 gRNA groups.

#### Distal enhancer positive control gRNA

15 gRNA targeting the *HBE1* TSS, and HS1-4 of the Globin LCR were chosen as validated from (*10*, *35*). These were paired based on target-site to create a gRNA-group.

#### Additional gRNA

We also included a small set of gRNA that targeted variants that were functional in massively parallel reporter assays (Ulirsch et al. 2016). However, as only intragenically targeting gRNA created a significant effect on its host gene in this dataset (and thus were potentially confounded by CRISPRi interfering with transcription via a steric rather than epigenetic mechanism), we did not report on these results here.

### 1,119 Candidate Enhancer At-Scale Library (1,119plex library, **Table S2**)

#### Picking candidate enhancer regions

K562 DNase-seq narrowPeaks (ENCSR000EKS) < 1 Kb away from any gene (GENCODE March 2017 v26lift37) were bedtools-intersected with K562 HiC domains (*24*) that contained at least one of the top 10% highest express genes in the 6,806 cell pilot study K562 data set. The remaining regions were tested for intersections with K562 GATA1 ChIP-seq narrowPeaks (ENCSR000EFT, lifted to hg19), H3K27ac ChIP-seq narrowPeaks (ENCSR000AKP, lifted to hg19), RNA Pol II ChIP-seq narrowPeaks (ENCSR000AKY), and EP300 ChIP-seq narrowPeaks (ENCSR000EHI) (**Fig. 3A**).

#### Candidate enhancer gRNA

NGG-protospacers within these candidate enhancers were again scored using default parameters of FlashFry (*40*), and the two top quality-scoring gRNA per region were chosen as spacers to be used in the gRNA library (scores prioritized by Doench2014OnTarget > Hsu2013 > Doench2016CDFScore > otCount). The same spacers from the 100plex library that were putative candidate enhancers of *NMU* and *HBG1* were also added.

#### TSS positive control gRNA

380 genes were randomly sampled from the highly-expressed genes within the same HiC domain (as described above) and 2 gRNA were again chosen per gene from spacers with the best empirical and predicted scores of the hCRISPRiv2 library (*39*). The gRNAs targeting the TSS of NMU was again included for a total of 381 TSS targeted.

#### NTC gRNA

All original NTC were again included from the 100plex study. More were chosen from 6 random regions of the hg19 genome (chr4:25697737-25700237, chr5:12539119-12541619, chr6:23837183-23839683, chr8:11072736-11075236, chr8:23768553-23771053, chr9:41022164-41024664) using FlashFry (*40*) to total 50 targeting these gene-devoid regions of the genome. A further 50 were derived from the 11 scrambled sequences used in the 100plex library and 39 NTC sampled from those recommended by (*39*).

#### Distal enhancer positive control gRNA

Also included were 14 of the gRNA from 100plex library that targeted the globin LCR.

Of note, our initial FlashFry quality annotations did not label a small number of protospacers with perfect repeat off-targets, permitting their inclusion in our library (81 of 2,238 spacers ordered in the 1,119plex library; only 9 gRNA-groups with both spacers affected). gRNA-groups with an impacted spacer were rare in our significant crisprQTLs. We also note that we still expect these guides to target their intended site.

## gRNA-library cloning, and transduction

The lentiviral CROP-seq gRNA-expression vector (*16*) was modified by Q5-Site Directed Mutagenesis (New England BioLabs, F:5-acagcatagcaagtttAAATAAGGCTAGTCCGTTATC-3 R:5-ttccagcatagctcttAAACAGAGACGTACAAAAAAG-3) to incorporate the previously described gRNA-(F+E)-combined backbone optimized for CRISPRi (*17*)(*18*). Prepared vector was digested with *BsmBI (*FastDigest Esp3I, Thermo Fisher Scientific), “filler” sequence removed by gel extraction, and cleaned (Zymo Research DNA Clean & Concentrator-5) vector without “filler” was used for all downstream cloning.

Protospacer libraries were ordered single stranded from Integrated DNA Technologies (100plex library: as separate oligos and then pooled) or as a pool (1,119plex library: CustomArray Inc., each protospacer synthesized in triplicate in the pool). Pools were amplified (5-atcttGTGGAAAGGACGAAACA-3, 5-acttgctaTGCTGTTTCCAGC-3, Kapa Biosystems HiFi Hotstart ReadyMix (KHF)) and purified amplicons (Zymo Research DNA Clean & Concentrator-5) were cloned into CRISPRi-optimized CROP-seq vector prepared as described above (NEBuilder^®^ HiFi DNA Assembly Cloning Kit, NEB). Products were transformed into Stable Competent *E. coli* (NEB C3040H) to produce >20 colonies per gRNA in the library.

The Fred Hutchinson Co-operative Center for Excellence in Hematology Vector Production core produced all virus. Cells were transduced with varying titers of virus to achieve differing MOI with 8 μg/mL polybrene. At 24 hours post-transduction, cells were spun and resuspended with virus-free media. At a total 48 hours post-transduction, 2 μg/mL puromycin was added to the culture, and changed to 1 μg/mL puromycin at the next passage for maintenance. A total of 10 days post transduction, cells were collected for scRNA-seq.

## Single cell RNA sequencing

~4000-8000 cells were captured per lane of a 10X Chromium device using 10X V2 Single Cell 3’ Solution reagents (10X Genomics, Inc). A single lane was used for the 100plex experiment, while six lanes were used for both the low and high MOI 1,119plex experiments. All protocols were performed as per manufacturer’s recommendations, except prior to the enzymatic shearing step, a portion of full length cDNA was taken for PCR enrichment of gRNA-sequences off the CRISPRi-optimized CROP-seq transcripts as described below.

Final libraries were sequenced on a NextSeq 500 with 75-cycle high-output kits (R1:26 I1:8 I2:0, R2:57). The 100plex library lane was sequenced to 24.1 thousand median UMI per cell, and 44.4% sequencing saturation. The low MOI 1,119plex library lanes were sequenced to 16.7, 20.9, 21.8, 22.7, 23.4, 25.6 thousand median UMI per cell, at 16-25% sequencing saturation. The high MOI 1,119plex library lanes were sequenced to 16.1, 18.5, 19.5, 19.8, 20.3, 20.6 thousand median UMI per cell, at 16-20% sequencing saturation. All quality statistics are reported from Cell Ranger defaults.

## gRNA-transcript enrichment PCR

A three-step hemi-nested PCR reaction was performed to enrich gRNA sequences from the 3’ UTR of puromycin resistance gene transcripts produced by the CRISPRi-optimized CROP-seq integrant. PCR was monitored by qPCR to avoid overamplification, and each reaction was stopped immediately before it reached saturation.

### PCR 1

10-13 ng of full-length 10x scRNA-seq cDNA were amplified in each 50 μl KHF reaction, spiked with SYBR-Gold (Invitrogen) for qPCR monitoring (3% of all cDNA for the 100plex experiment versus 10% of all cDNA for 1,119-plex experiments). Greater input appeared to lead to less cells with incomplete gRNA assignments, as indicated by **Fig. 3B** versus the more zero-inflated distribution of **Fig. S1A**.

F: U6_OUTER 5- TTTCCCATGATTCCTTCATATTTGC -3

R: R1_PCR1 5- ACACTCTTTCCCTACACGACG-3

PCR 2: 4% of each previous reaction was brought forward, sample replicates pooled, cleaned with 1x Agencourt AMPure XP beads (Beckman Coulter), and 1/25th of the cleaned pooled product was amplified in each of six 50 μl reactions.

F: U6_INNER_with_P7_adapter

5-GTCTCGTGGGCTCGGAGATGTGTATAAGAGACAGcTTGTGGAAAGGACGAAACAC-3

R: R1-P5 5-AATGATACGGCGACCACCGAGATCTACACTCTTTCCCTACACGACG-3

PCR 3: The PCR 2 replicate reactions were pooled, 1x AMPure cleaned, and 1/25th of the cleaned pooled product was amplified in each of six 50 μl reactions and products cleaned once again via 1x Ampure.

F: 5- CAAGCAGAAGACGGCATACGAGATIIIIIIIIIIiGTCTCGTGGGCTCGG-3 (standard NEXTERA P7 indexing primer)

R: R1-P5 again

## Digital gene expression quantification

Sequencing data from each sample was processed using the Cell Ranger software package as provided by 10x Genomics, Inc., to generate sparse matrices of UMI counts for each gene across all cells in the experiment.

Each lane of cells was processed independently using cellranger count, aggregating data from multiple sequencing runs. The 100plex library experiment was processed with cellranger 1.3.1 and the 1,119plex library experiments were each processed with cellranger 2.0.2.

### Definition of genes well-expressed or ‘detectably expressed’ in K562

Unless otherwise notes, genes were defined as well expressed or detectably expressed in K562 if they had at least one read in 0.525% of cells (250 cells) in the high-MOI 47,650-cell dataset (12,984 total genes).

### Assigning genotypes to cells

gRNAs were assigned to cells as performed in Hill, McFaline-Figueroa et al. (*18*); briefly, sequences corresponding to the gRNA-containing CRISPRi-optimized CROP-seq transcripts are extracted from the cellranger position sorted BAM file after running our custom indexed libraries through the cellranger pipeline to tag reads with corrected cell barcodes and UMIs. gRNA sequences are extracted and corrected to the library whitelist within an edit distance of two,and gRNA-cell pairs are tracked when a valid cell barcode and UMI are both assigned to the read. Likely chimeric reads are detected and removed to reduce noise in the assignments as previously described. As in Hill, McFaline-Figueroa *et al. (18)*, we utilized thresholds to set minimum acceptable values for the total reads for a gRNA-cell pair and for the proportion of all CROP-seq transcript reads accounted for by each gRNA observed in a cell to distinguish noise from real assignments. Here, given the larger number of guides contained in each cell, we find that UMI counts provide a much cleaner distribution than read counts and have used UMI counts in all calculations. For the 100plex library, we used cutoffs of 0.035 and 5 UMIs as the thresholds for the fraction of CRISPRi-optimized CROP-seq transcripts attributed to a guide and the total number of observed UMIs for a gRNA-cell pair, respectively. For the 1,119plex library experiments we used 0.01 and 5 UMI in both our low and high MOI for each of these thresholds. Only cell barcodes that appear in the set of passing cells output by cellranger, which imposes an automated threshold on the total UMIs observed in cells, are carried forward in downstream analysis.

### Differential expression tests

In our *cis* analyses, we tested each perturbing gRNA-group against a gene within 1 Mb of the gRNA. These gRNA-gene pairs were identified by using bedtools to intersect the DHSs targeted by the gRNA library with 1 Mb windows in either direction of TSS annotations from GENCODE March 2017 v26lift37 (total of 2 Mb, centered around the TSS). In our *trans* analysis, all gRNA-groups were paired with all genes that were defined as expressed in K562. In both *cis* and *trans* analyses, NTCs were tested against any genes used to test perturbing-gRNAs.

For each gRNA-group we assigned a label of “1” to cells that contained a gRNA belonging to that group and a label of “0” to all other cells in the dataset. Monocle2 was used to perform a differential expression test, using the negative binomial family, over this categorical label to find differentially expressed genes between these two groups. Due to its support of complex model formulas, Monocle2 does not provide model coefficients as part of the differential expression results. We created a modified version of the differentialGeneTest function and associated helper functions that return both the intercept term and the coefficient of the group assignment to facilitate more robust prioritization and characterization of hits from our screen. The negative binomial family uses log as the link-function, so we can calculate the initial expression level as exp(intercept), and the fold change in expression between the two groups as exp(group_coefficient + intercept) / exp(intercept). We verified data from our power simulations that the appropriate effect sizes can be obtained with this method using the coefficients output by VGAM.

## Calling hits from differential expression test results

In both the *cis* and *trans* analyses, all differential expression test results were filtered to only K562 expressed genes.

Tests with two sources of potential false positives were excluded:

1. We identified inflation of NTCs when testing them against genes highly impacted by perturbing-gRNA in our library (for example, NTCs associated with targets of our TSS and globin LCR controls). This was due to subtle yet detectable nonrandom associations of gRNA-groups with other gRNA-groups across cells. To exclude this source of inflation, we used Fischer’s exact test to identify when an NTC was nonrandomly assorted with a perturbing-gRNA (*q-*value < 0.01 & odds ratio > 1). Then, any test of an NTC against a gene within 1 Mb of that gRNA’s gRNA-group was excluded from further analytical steps. This was used to exclude such interactions in both the *cis* and *trans* analyses.
2. We noted Monocle was susceptible to inflating *P-*values when a gene was highly expressed but only in few cells. To avoid this problem, we excluded 23 outlier genes that were expressed in < 20,000 cells in the high-MOI 47,650-cell dataset and with log10(total reads / cells with a read) >0.2 greater than predicted by a spline fit generated via smooth.spline() with spar=0.85 to limit overfitting (**Fig. S9**).

Remaining tests were filtered to those that decreased expression of the target gene. Then, a 10% empirical FDR was defined as the *P*-value at which the proportion of passing NTC-tests/total NTC-tests was 10% of the proportion of passing candidate enhancer tests/total candidate enhancer tests. *cis* pilot *P-*value = 5.52e-11; *cis* high MOI *P-*value = 5.82e-05; *trans* high MOI *P-*value = 8.45e-13; *cis* low MOI *P-*value = 7.87e-05.

## Aggregate analysis of crisprQTLs

### Distance between enhancer and eCRISPR-gene

Distance was calculated between the GENCODE March 2017 v26lift37 annotated TSS of the perturbed gene and the leftmost gRNA (if targeting a candidate enhancer) or the GENCODE-annotated TSS of the originally targeted transcript (if targeting a TSS).

### ChIP-seq strength quintile analysis and logistic regression classifier

1,119 candidate enhancers were bedtools-intersected with ChIP-seq of histone-associated marks (all narrowPeak; CCNT2 ENCSR000DOA, H3K27ac ENCSR000AKP, ENCSR038DJJ SMAD1, H3K4me1 ENCSR000AKS, RNA Pol II ENCSR388QZF, TAL1 ENCSR106FRG, EP300 ENCSR000EGE, ENCSR000EFT GATA1, ENCSR000EWG GATA2 (*1*)), broken into quintiles of the 7th “signalValue” column (representing peak strength, usually representing overall average enrichment in the region), and the rates of crisprQTL identified in each quintile were used. The signalValue column of each dataset was also used to fit both independent and multivariate logistic regression classifiers using the glm() function with binomial family. Fivefold cross-validation was

### Functional annotation enrichment

We used the Piano package (*32*) to perform functional annotation enrichment from the ‘all pathways’ Gene Ontology (http://download.baderlab.org/EM_Genesets/June_20_2014/Human/June_20_2014_versions.txt).

In the *cis* analysis, the 5,751 K562-expressed genes within 1 Mb of a perturbing-gRNA were used as our background dataset, and 105 eCRISPR-genes randomly sampled from genes with expression greater than the mean of our 105 eCRISPR-genes was used as the comparison set of “expression matched controls”. In the *trans* analysis, K562 expressed genes were used as a background. In both analyses, the most significant terms were described in the text.

## Replication of crisprQTL outside of multiplexed experiment

To replicate the CRISPRi phenotype outside of the single-cell pooled screen, we prepared small pools of gRNAs targeting e-NMU (hg19 chr4:56595855-56597095), e-PRKCB (chr16:23833191-23833694), e-PTGER3 (chr1:71140581-71141997), e-GYPC (chr2:127402697-127403360), e-HMGA1 (chr6:34191315-34192504), or the TSSs of their respective eCRISPR target genes (**Table S6**). These small gRNA pools were cloned into CRISPRi-optimized-CROP-seq (as described above). Lentiviral preps from these gRNA pools were transduced at low MOI into the K562-dCas9-BFP-KRAB line, and cultured for 10 days under puromycin selection before two replicates of RNA were collected. Bulk RNA-seq libraries were prepared from each replicate via a TruSeq RNA kit (400 ng input), and sequenced on a NextSeq 500 with two 150-cycle kits in mid output mode. Gene-level quantifications and differential expression tests were performed via kallisto (*41*) and sleuth (*42*). The quantification uncertainty for each sample was estimated by the kallisto and sleuth bootstrapping procedure (n=100 bootstraps).

## Power Simulations

In order to predict the impact of multiplexing on the power of crisprQTL experiments, we developed a simulation framework. First, using single-cell RNA-seq data collected from the 1,119plex 47,650 K562 cells, we estimated a dispersion function that relates the mean expression of a gene to its dispersion estimate (one of the two parameters required for the negative binomial distribution) calling the Monocle2 functions estimateSizeFactors and estimateDispersions. This function is typically used in differential expression testing to shrink dispersion estimates, but here we use it to estimate dispersion values for simulated transcripts. This dispersion function is then extracted from the CellDataset object output by Monocle2 and used as input to our simulations.

Next, we chose relevant ranges for each of the parameters varied in our simulation: the MOI, total cell count, effect size (fraction repressed by CRISPRi), and mean expression level of the gene being tested. By examining the range of expression values observed in our data, we chose to simulate expression data for genes having mean expression values (size parameter of the negative binomial distribution) of 0.01, 0.1, 0.32, 1.0, 3.16, and 10.0 UMIs (0.10, 0.32 and 1.00 used respectively as low, medium, and high in **Fig. 1B**) to provide a range of representative values.

We simulated MOIs at several values from 0.3 to 50, a range which includes the MOIs estimated from our own crisprQTL mapping screens. For each MOI, we calculate the expected number of cells containing a given guide by assuming a Poisson distribution of lentiviral delivery, zero-truncating the distribution to account for drug selection for cells that contain a guide transcript, and rescaling the probability distribution of guide counts accordingly. Perfect library uniformity was assumed to obtain the expected number of cells containing a given guide and the number of cells that do not contain that guide. Effect sizes of CRISPRi repression were chosen using estimates from the literature and were simulated at several values between 10% to 90% percent repression of the average expression level of the target transcript (size parameter input to the negative binomial distribution).

Finally, we simulated several values of total cells included in the experiment ranging from 35,000 to 300,000 cells (45,000 cells shown in **Fig. 1A**). Expression data from transcripts corresponding to 100 samplings per set of parameters were generated for the populations of cells containing the gRNA and not containing the gRNA respectively. Our expression data simulation assumed a negative binomial distribution with the appropriate size parameter for the cells with and without the gRNA, and a dispersion value estimated using the dispersion function described above given the starting mean expression level being simulated. For each set of parameters, the simulated transcripts were subjected to a differential expression test performed between cells with and without the gRNA assigned using our modified version of the Monocle2 function differentialGeneTest as described above (see Differential Expression Tests). *P-*values were obtained and corrected assuming an average number of 20 tests per group in the library to approximate the number of genes contained within 1 Mb on either side of each gRNA-group and the impact of multiple testing. The rate of tests falling below a *q-*value of 0.05 were tabulated at each set of parameters to make power curves.

**Fig. S1.**
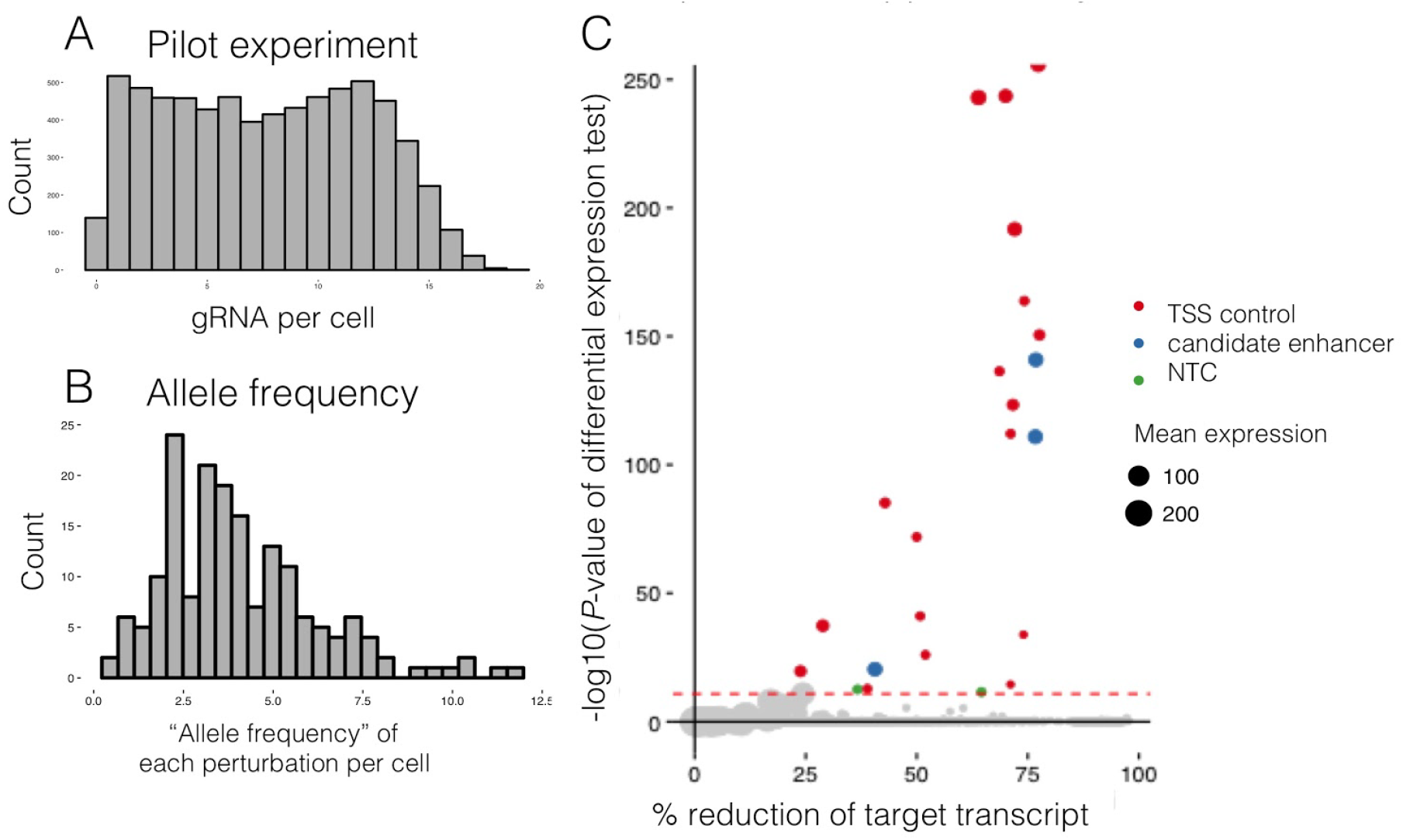
Extended data for 100 candidate enhancer pilot experiment. (A) Number of gRNAs identified per cell. Median 8 gRNA +/- 4.5 gRNA were identified per cell from targeted amplification of the pre-fragmentation 10x cDNA library. However, the amount of cDNA input used for this amplification was less than used in later experiments. **(B)** “Allele frequency” of each perturbation in the cell population, *i.e.* the number of cells in which a given site was targeted. Mean 4.1% +/- 2.2%. Sites targeted by >2 gRNAs not included here. **(C)** Distribution of all differential expression tests (significance x effective size) that decrease test-gene expression. log10(*q*-val) versus % the target transcript is reduced. Point size corresponds to relative expression level. Dotted red line = *P-*value threshold used to call hits.

**Fig. S2.**
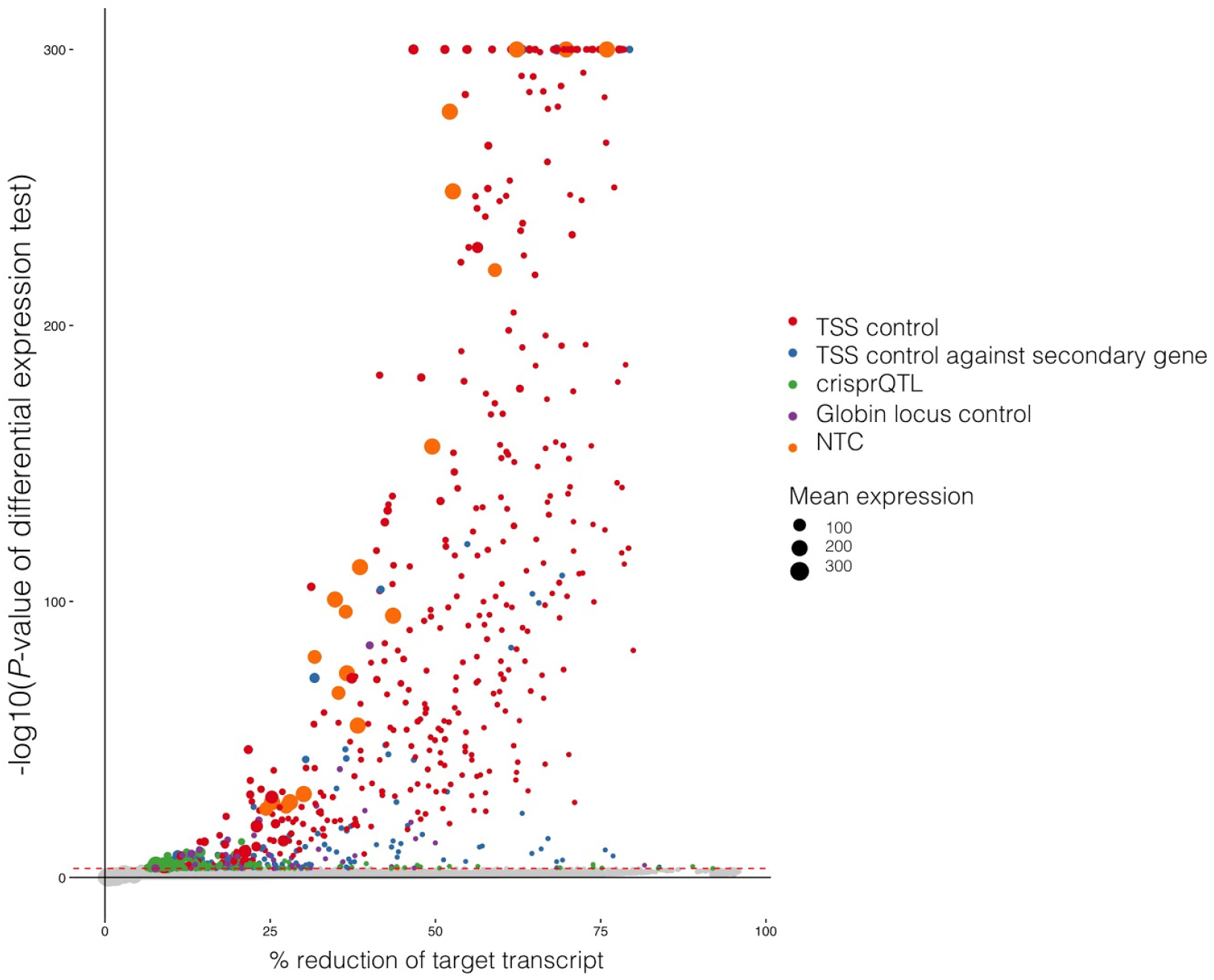
Distribution of all differential expression tests for the 1,119 candidate enhancer crisprQTL experiment. Effect size vs. significance is plotted for all tests that showed a decrease in test gene expression. log10(*P-*value) versus % target transcript is reduced. Point size corresponds to relative expression level. Dotted red line = *P-*value threshold used to call hits.

**Fig. S3.**
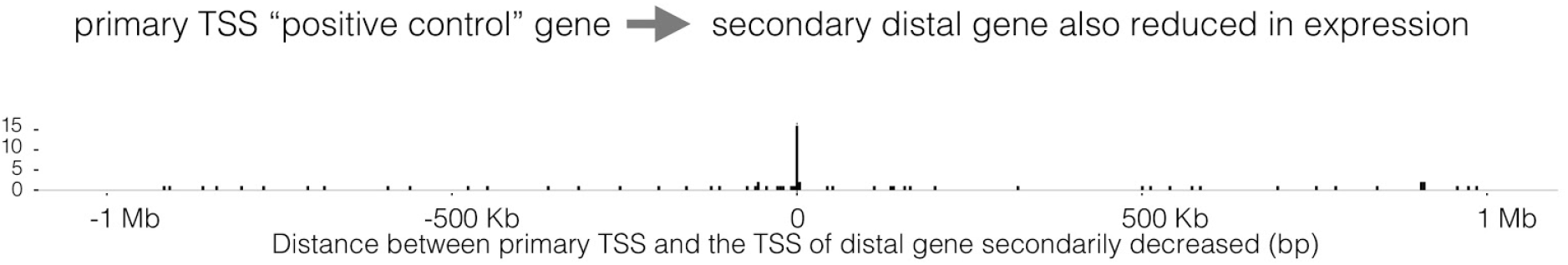
Distribution of distances between “positive control” TSSs and secondarily repressed genes. Distance between the 56 “positive control” targeted TSSs and the TSS of the secondary gene (over 1 Kb away) whose expression was simultaneously also decreased. Plotted with respect to primary target gene’s orientation.

**Fig. S4.**
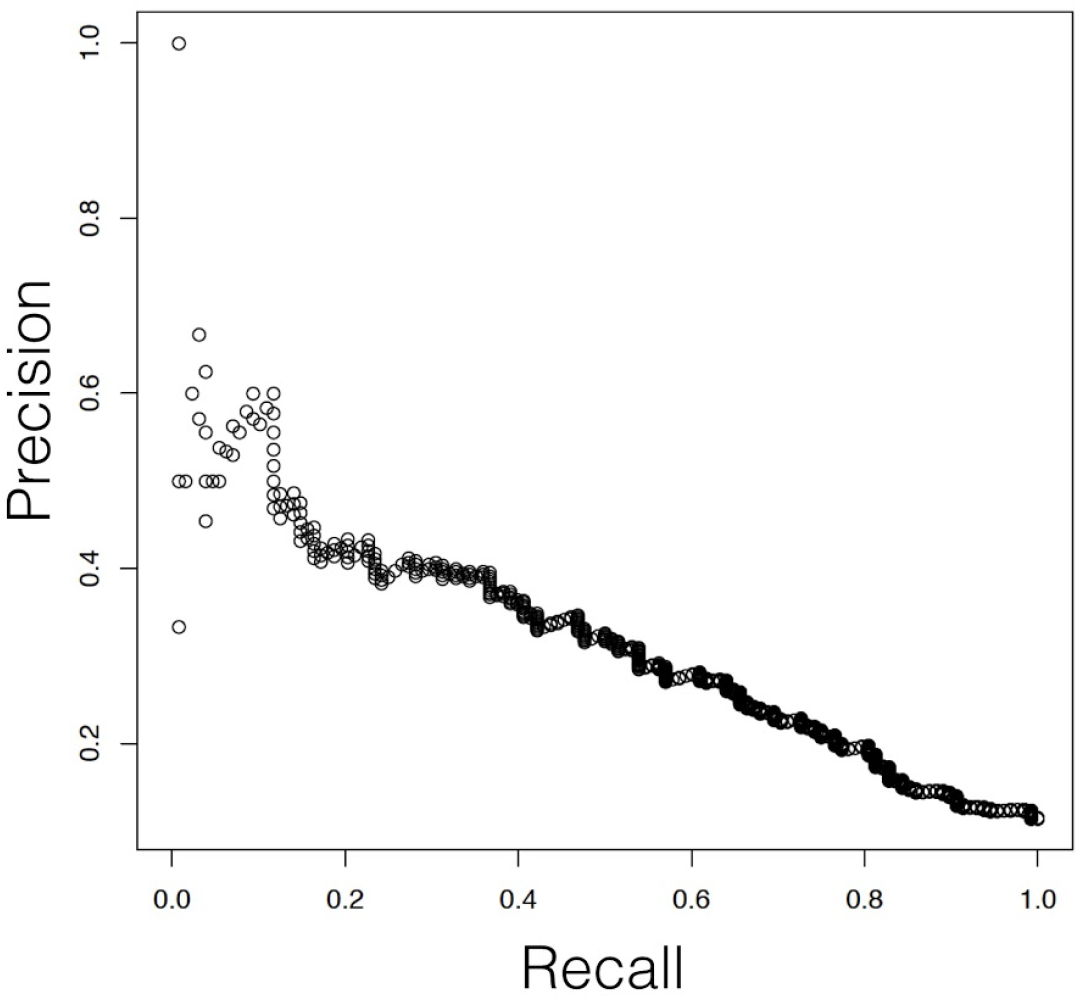
Precision recall curve for a multivariate logistic regression classifier based on ENCODE enhancer-associated biochemical features that differentiates experimentally identified crisprQTLs from the 1,119 candidate enhancer library. The median AUPR from fivefold cross-validation was 0.31.

**Fig. S5.**
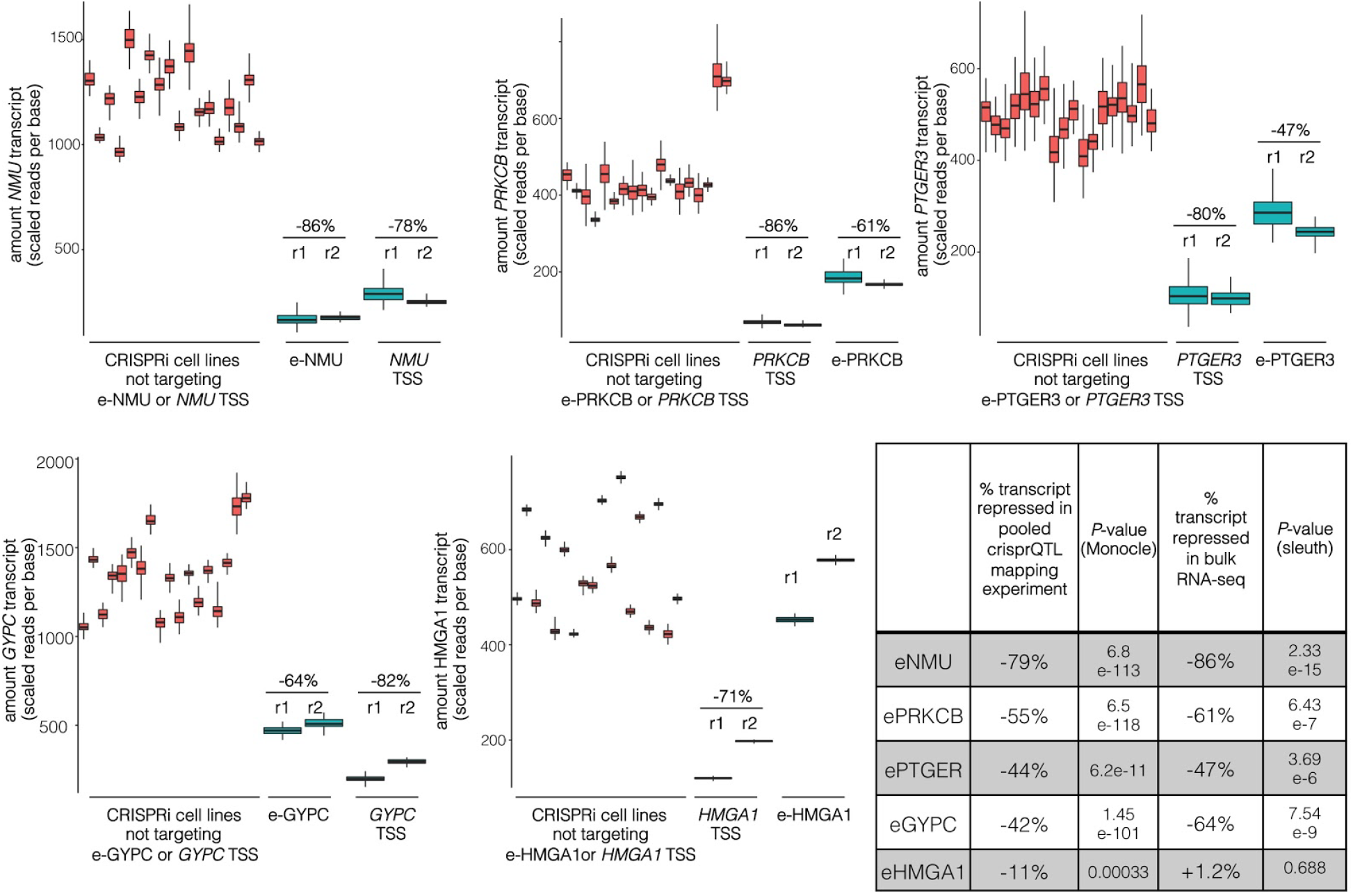
Replication of crisprQTL outside of multiplexed mapping experiment. Two replicates of bulk RNA-seq were performed on RNA extracted from CRISPRi-positive K562 separately transduced with sgOPTI-CROP-seq gRNAs solely targeting e-NMU (hg19 chr4:56595855-56597095), e-PRKCB (chr16:23833191-23833694), e-PTGER3 (chr1:71140581-71141997), e-GYPC (chr2:127402697-127403360), e-HMGA1 (chr6:34191315-34192504), or the TSSs of their respective eCRISPR target genes. Error bars represent the quantification uncertainty for each sample as estimated through the bootstrapping procedure implemented in kallisto and sleuth. Percent repression is calculated from Monocle differential expression test (in the case of the crisprQTL mapping experiment) or from the ratio of: (mean transcript in targeted RNA) / (mean transcript in non-targeted RNA) (in the case of bulk RNA-seq, as called by sleuth and kallisto).

**Fig. S6.**
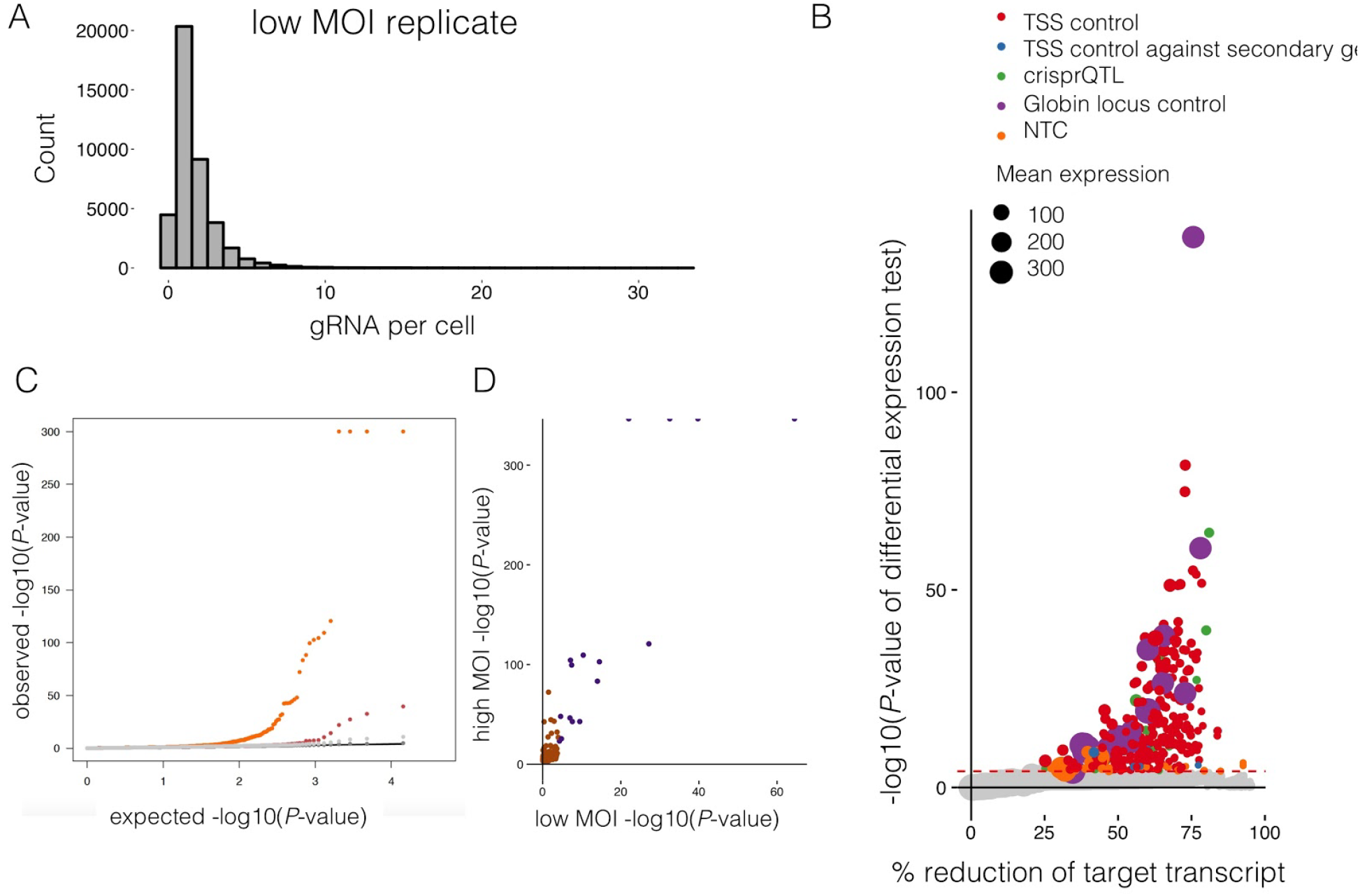
Extended data of the low MOI biological replicate. (A) gRNA/cell. The 1,119 K562-candidate enhancer library was transduced at an MOI of ~1. **(B),** Differential expression tests results for the low MOI experiment. Colored points correspond to perturbations called as ‘hits’ in this study, dashed red line shows the *P*-value threshold that corresponded to an empirical FDR of 10%. **(C),** Q-Q plot of the low (red) versus high (orange) MOI experiments shows an increase in power in the high MOI experiment, as seen by further inflated *P-*values. **(D),** *P*-values from the low versus high MOI studies. Only crisprQTLs found to be significant in the high MOI study are plotted (Pearson’s r = 0.334, purple = hit in both, orange = hit in just the high MOI study).

**Fig. S7.**
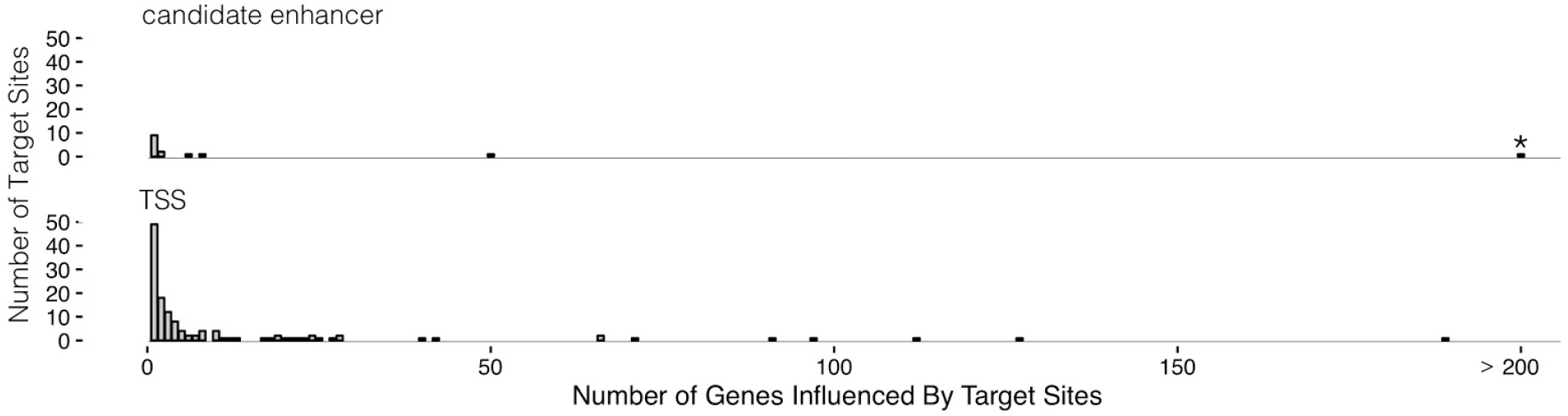
Extended data for *trans* crisprQTLs. Histogram of the number of eCRISPR-genes significantly impacted by *trans* crisprQTLs. (^*^ = one candidate enhancer located 75 Kb upstream of *LMO2* was found to act as a *trans* crisprQTL of 1,177 eCRISPR genes).

**Fig. S8.**
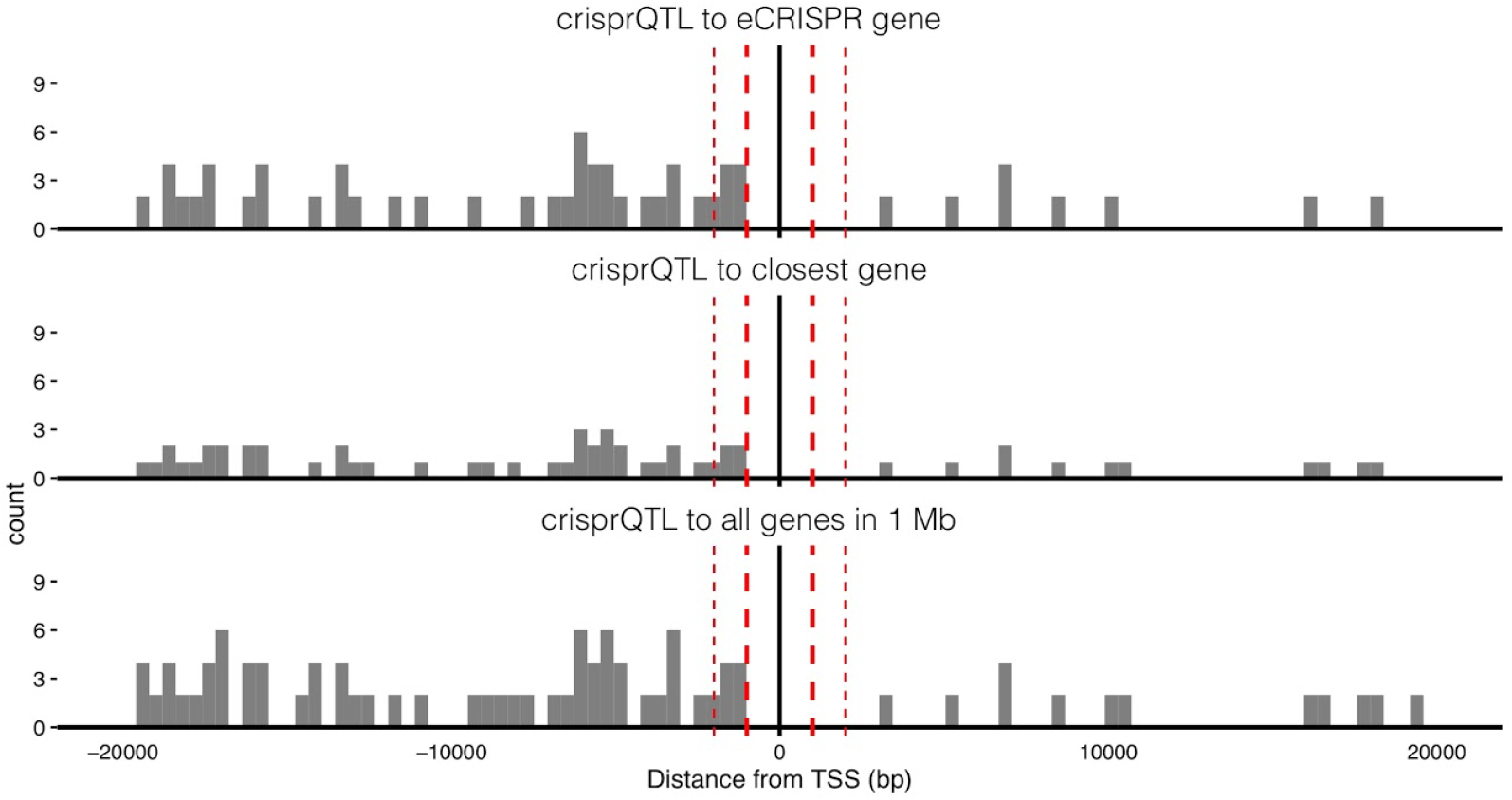
+/- 20 Kb zoomed in view of distance distribution between target gene and targeted site. To note, gene bodies were excluded from targeting to avoid potential CRISPRi interference with transcription. We also excluded any site within 1 Kb upstream of TSSs and 1 Kb downstream of TESs. (Thin dotted red line = 2 Kb from TSS, thick dotted red line = 1 Kb from TSS, black line = TSS)

**Fig. S9.**
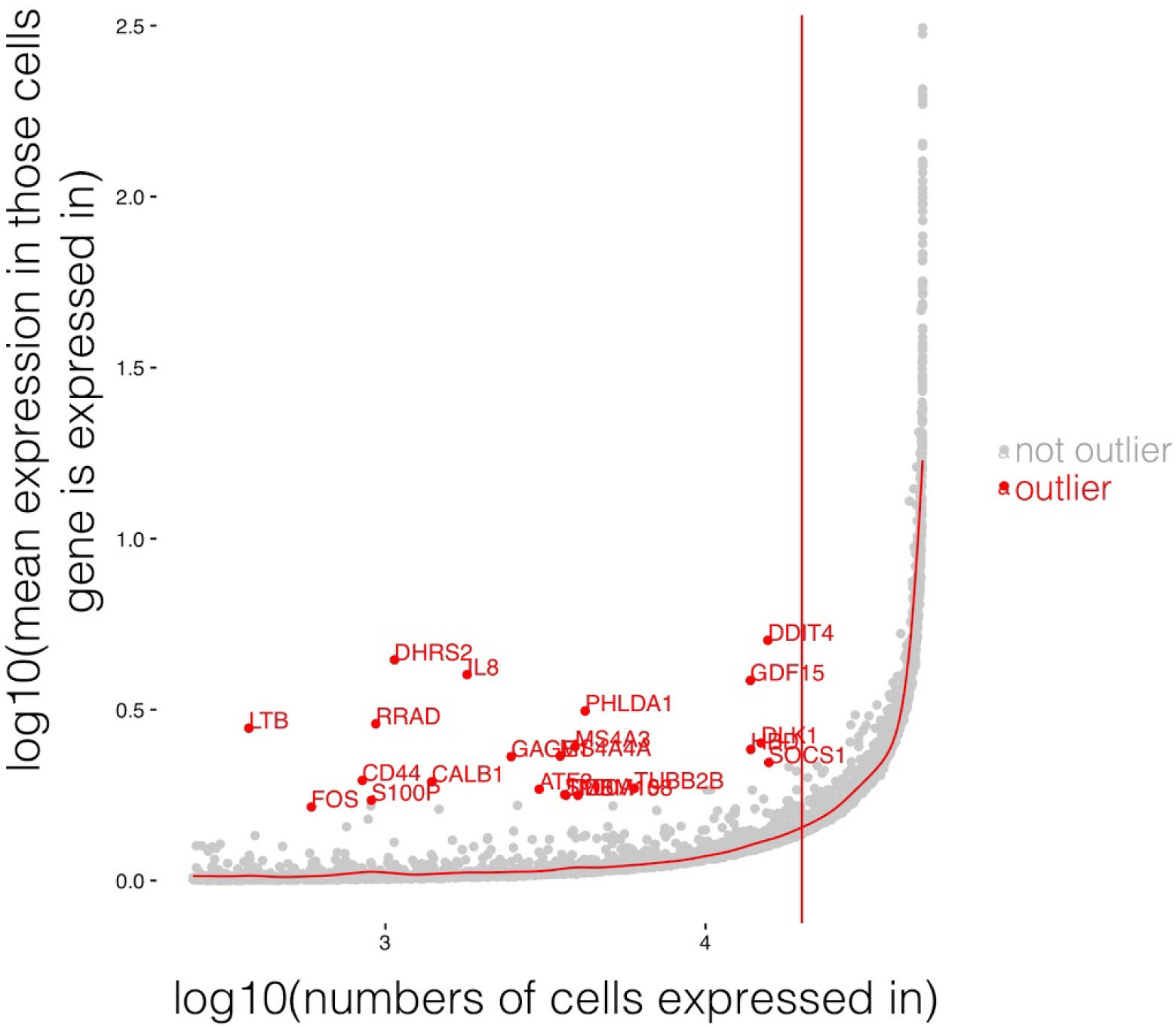
Excluded 23 expression outliers. We observed that differential expression test *P-*values were inflated when a transcript was highly expressed but only in few cells. Thus, we excluded 23 outlier genes (red) that were expressed in fewer than 20,000 cells in the high-MOI 47,650-cell dataset that also had >0.2 greater log10 (mean expression in those cells gene is expressed in) than predicted.

**Table S1 | 384 oligos used in pilot 100plex-library experiment.**

**Table S2 | 3,118 oligos used in scaled-up 1,119plex-library experiment.**

**Table S3 | Supplementary information of 145 cis crisprQTLs.**

**Table S4 | Ontological enrichment results of 105 eCRISPR genes.**

**Table S5 | Logistic regression P-values for independent and multivariate models.**

**Table S6 | Oligos used for replication of selected crisprQTLs outside of pool.**

